# Justify your disability! A simulated medical evaluation as a robust novel stress induction paradigm in chronic pain patients

**DOI:** 10.1101/2024.07.03.601919

**Authors:** Aimie Chiron, Aurore Fernandez, Léa Schmidt, Sonia Plaza-Wüthrich, Marc R. Suter, Siri Leknes, Chantal Berna

## Abstract

Maladaptive stress responses may exacerbate chronic widespread pain (CWP) and deserve further investigations. Yet, existing stress induction paradigms lack relevance for individuals with this condition. Hence, we developed the Social Benefits Stress Test (SBST), adapted from the Trier Social Stress Test. Instead of a job interview, the main task consists in justifying the inability to work.

Forty women with CWP in the context of hypermobile Ehlers-Danlos syndrome or hypermobility spectrum disorders were included. They underwent a 30-min baseline, the new stress task and a recovery period. The psychophysiological stress response was captured using self-reported stress ratings, salivary cortisol and α- amylase levels, as well as continuous physiological monitoring of heart rate variability (HRV) and electrodermal activity (EDA).

Compared to baseline, the analysis revealed a significant and transient increase in stress ratings during the stress task, associated with a peak in salivary biomarkers concentrations. The HRV signal analysis showed a significant decrease in high frequency power (HF), and increases in heart rate, low frequency power (LF) and in LF/HF ratio. The EDA analysis revealed a significant increase in skin conductance level (SCL) tonic component and skin conductance response (SCR). Subjective stress ratings positively correlated with changes in salivary biomarkers, LF/HF ratio and EDA outcomes.

The SBST induced a reproducible moderate stress response across subjective and physiological measures in a population of CWP patients, validating this task as a relevant experimental model of social stress in chronic pain. The SBST is a useful tool to study the relationship between stress and chronic pain.

**Perspective:** This manuscript presents the Social Benefits Stress Test (SBST) as a novel paradigm to assess stress reactivity in chronic widespread pain patients. By simulating the challenge of justifying work incapacity, it elicits a reproducible stress response, supporting its use as a model to study stress-pain interactions and evaluate therapeutic interventions.

## Introduction

Chronic pain affects around 20% people worldwide, representing a physical and socioeconomic burden^35,58^, where one of the main issues is an inability to work^37,80^. However, functional impairment is often met by disbelief in medico-social systems^27,60,72^ due to the disease invisible nature^20,90^. These issues are key to livelihood and sense of identity of patients with chronic pain, and represent major stressors^59,60^.

Patients’ stress elicited by the denial of support or others’ doubts of the illness is under-investigated^90^. Yet there is evidence that stigma, social support (or lack thereof), and interactions with healthcare and social services are key factors shaping their well-being^2,6,52,59,60,66,84^. Socioeconomic struggles could also contribute to the vicious cycle of chronic pain. In fact, prolonged exposure to stressors can result in long-term maladaptive stress responses, which are associated with worsening^82,83^ and maintenance of chronic pain^9,36,81,86^. Chronic stress involves a dysregulation of the neuro-endocrine cascade of the hypothalamo-pituitary-adrenal (HPA) axis and the autonomous nervous system (ANS) balance^31^. It appears these interactions are specifically relevant for the pathophysiology of chronic widespread pain (CWP)^56,73,87^.

Induced acute stress responses in chronic pain have mainly been studied using the gold standard social stress induction task, the Trier Social Stress Test (TSST)^12,23,41,85^. However, due to its reliance on a mock job interview, the TSST may not be effective in this population: the disconnect between the test scenario and their real-life experiences could result in disengagement or overwhelming distress. Results may be further muddled by an elevated baseline stress level and an increased vulnerability to social stress^23^. The TSST has previously been adapted in other contexts where a job interview seemed irrelevant^39,70,88,89^. Here, we validate the Social Benefits Stress Test (SBST), a TSST adaptation tailored to people suffering from disabling chronic pain, created as a research tool for future work in the interconnecting fields of chronic pain and stress. The job interview was replaced by a simulated interview with an expert deciding on patients’ work incapacity for social benefits allocation. The main measures were subjective stress outcomes on visual analogue scales (VAS), heart rate variability (HRV), electrodermal activity (EDA), as well as salivary cortisol and α-amylase. To be considered as a valid stress induction task, the SBST had to induce a significant and transient increase in subjective stress levels, associated with congruent increased cortisol, α-amylase levels, decreased HRV and increased EDA.

The patients were suffering from hypermobile Ehlers-Danlos Syndrome (hEDS) and Hypermobility Spectrum Disorders (HSD). hEDS/HSD are incapacitating generalized connective tissue disorders with joint hypermobility often presenting with CWP, musculoskeletal complications and evidence of small fiber neuropathy^32,48^. Many suffer from fatigue, sleep disorders, anxiety, and depression^13^. The disease load is often incompatible with full-time employment^8^. Yet, being an invisible, late-diagnosed, poorly recognized condition, there is a frequent lack of knowledge and recognition from health care and social support providers^30^. Hence, exploring subjective and objective stress parameters in response to a social stress induction task in this population seemed particularly relevant.

## Methods

### Participants

This prospective single-center study took place at the Centre for integrative and complementary medicine at the Lausanne University Hospital. This study was conducted in accordance with the Declaration of Helsinki and was approved by the local ethics committee for research in humans (CER Vaud - 2021-01750). Participants were thanked for their participation with a 50 CHF voucher for a local store and their travel expenses were reimbursed.

Eligible participants were adult females, suffering from hEDS or HSD according to Malfait criteria^17,48^ reporting chronic pain (more than 6 months), willing and able to give informed consent, recruited from the Pain centre and Centre for integrative and complementary medicine. Since 89% of the patients suffering from hEDS/HSD are women^16^ and to avoid the confounding effect of sex, as it may affect stress responses, only women were included. Exclusion criteria were insufficient French language skills to communicate without a translator, severe hearing impairment, unstable psychiatric co-morbidity, active substance use disorder, somatic co-morbidity that could interfere with interpretation of heart rate variability (HRV) recordings (including cardiac malformation, recent myocardial dysfunction, cardiac transplantation, or cardiac treatment i.e. pacemaker, antiarrhythmic drugs, muscarinic receptor blockers, angiotensin-converting enzyme (ACE) inhibitors or any that would induce a modification in heart rate)^65^ or contraindication to sensor positioning (i.e. local skin damage or allergies).

### The Social Benefits Stress Test (SBST)

#### Task development

The task was an adaptation of the TSST^41^, maintaining the global timeline and procedures, including filming the test and the surprise arithmetic task (Figure 1). The fake job interview was replaced by a simulated interview with a medical expert. Two patient-advocates, suffering from chronic pain for more than five years and with previous experiences with the social benefits system in Switzerland, were involved in the task design through semi-structured interviews. They provided advice on the medical expert’s attitude and specific questions.

**Figure 1:**
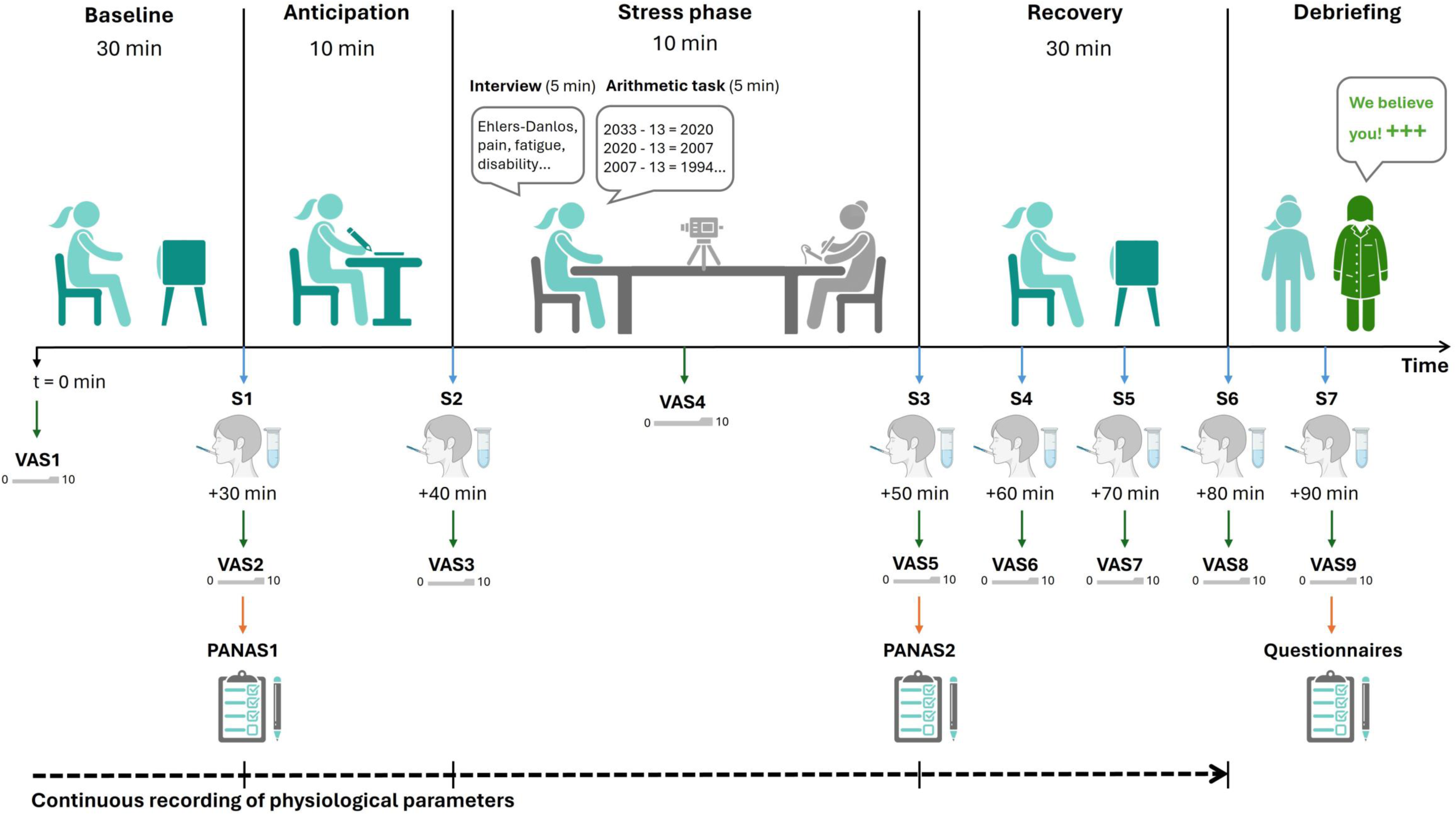
Design of the Social Benefits Stress Test (SBST). Participants underwent baseline, anticipation, stress, and recovery phases followed by a debriefing. Salivary samples (S1-S7) and stress and pain Visual Analog Scales (VAS1-VAS9) were collected at several timepoints. Positive Affects and Negative Affects (PANAS) questionnaires were filled before and after the stress phase. Physiological parameters were recorded continuously during the task

#### Procedure

Patients were invited for a unique study visit about stress (without further information regarding the task) lasting about 2h30, starting at 2PM for each participant. After collecting informed consent, patients received instructions for the saliva collection and sensors were positioned for ECG recording.

The task started with a 30min “**baseline**” with the participants siting on a comfortable chair and invited to relax while watching a marine life documentary (Océans, Jacques Perrin, 2009) (Figure 1).

After this baseline, they received the following instructions: “*Imagine* that you are not able to work full-time due to your pain, and you have asked for financial social support from an insurance company or the state. You will face a medical expert who will evaluate your work capacity. You now have 10 minutes to prepare for this interview.” Participants were also told that nonverbal behaviour and global performance would be analysed based on a video-recording.

This initiated the 10-minute “**anticipation**” phase, where patients were provided with a paper and pencil to prepare. The “**stress task**” followed: They were guided to a conference room, where they were introduced to the simulated medical expert. The medical expert was played by a female research collaborator (S.P.W.), in her forties, looking formal, severe, and sharp, dressed in a medical white coat. Her attitude was a “poker face”, imitating in the cold neutrality of the panel in the TSST, as well as attitudes of real medical experts described by patients advocates. She explained in a cold and distant way “You now have 5 minutes to justify your inability to work”. The interactions followed a protocol with scripted questions: If the subject spoke for less than 3 min, she said ‘You still have some time left. Please continue.’ If the subjects finished a second time before the 3 min were over, the expert stayed quiet for 20 s. In any case, after 3min, the expert asked prepared questions (see supplementary) during 2 min until the end of the 5 minutes. Following this “i**nterview**”, participants were asked to undergo a mental “**arithmetic task**” for 5 min, i.e. sequentially subtracting 13 from the number 2033 (as in the TSST).

Then the “**recovery**” phase followed: participants were guided back to the first room. They were invited to sit and relax while watching another 30 minutes of the marine documentary. After 30 min, the sensors were removed, and the patients were fully debriefed about the purpose of the task (“**debriefing**”). The research collaborator made it a point of honour to emphasize the recognition of their condition and provided a detailed clarification of the expected impacts of the study.

## Measures

### Subjective stress and pain

Ratings on visual analogue scales (VAS) were used to repeatedly assess subjective stress ^5^ and levels of pain ^64^ with two 10 cm-lines anchored “no <stress // pain>” on the left (= 0/10) and “worst <stress // pain> imaginable” on the right (= 10/10). Ratings were collected upon arrival and after each phase of the SBST (see Figure 1).

### Physiological recordings

Continuous physiological monitoring was performed using a 3-electrodes electrocardiogram (at a sampling rate of 1024 SPS) (NeXus-10 MKII, Mindmedia, Herten, the Netherlands). The physiological signals were recorded using the BioTrace+ software (Mind Media BV, Netherland).

Physiological data were analysed using the Neurokit2 Python library (v.0.2.3) ^47^. Initially, the raw data were exported, visually inspected, and corrected for artifacts and peak detection. Then HRV (root mean square successive differences (RMSSD), low-frequency power (LF) and high-frequency power (HF) components) ^14^ and EDA outcomes (*skin conductance response* (SCR) i-e the number of peaks for the phasic component and *skin conductance level* (SCL) for the tonic component ^10^) were calculated during designated 5-minute periods. For the phases lasting > 5 min, the extracted time-series were selected as a 5 min block outside of events of interference (e.g., sneezing, moving, talking, which had been noted with their timing during the testing). SCR peaks with a minimum amplitude of 0.1 µS were identified as events ^10^. This process was repeated for each phase of the task (baseline= 30 min, anticipation = 10 min, interview = 5 min, arithmetic= 5 min, recovery = 30 min phases). A typical stress response is characterized by a decreased HRV (with a decrease in RMSSD, an increase in LF, a decrease in HF, or an increase in the LF/HF ratio) and by an increased EDA (with an increase in the number of peaks and of the SCL ^11^) during the stress task compared to baseline recordings ^71^.

### Salivary cortisol and **α**-amylase

Saliva samples were collected using Salivettes (Sarstedt) to analyse the levels of salivary free cortisol and salivary α-amylase at the specified time points (Figure 1).

The participants were given instructions to abstain from eating, drinking, brushing their teeth, or smoking for 1 hour prior to the study visit. On the sampling day, they were also required to fill out a form, providing information about any medications or specific events that could potentially affect the analysis of collected biomarkers and none of them lead to data exclusion. Then, samples were centrifuged at +4°C (3’500 rpm, 15 min) and stored at −80°C until the analysis. Cortisol and α-amylase levels assessments were performed at the Psychological Institute laboratory of the University of Zürich. Cortisol levels (nmol/L) were assessed using a commercial luminescence immunoassay kit (ELISA; IBL International, RE62111/RE62119). For α- amylase analysis, saliva samples were diluted at a ratio of 1:400 with a NaCl solution and incubated with a substrate reagent, α-Amylase CC FS from Diasys, at 37°C. The increase in absorbance was transformed into α-amylase concentrations (U/ml) using a standard curve. Each sample was run in triplicate. Intra- and inter-assay coefficients of variation were reported to be below 10%.

### Questionnaires

Participants were asked to fill standardized questionnaires: the Brief Pain Inventory (BPI), assessing the pain severity and its impact on daily functions ^78^; the Perceived Stress Scale (PSS), evaluating how much their life was overwhelming and unpredictable over the past month ^21^; the Pain Catastrophizing Scale (PCS) quantifying rumination, magnification and helplessness regarding pain ^77^ and the Hospital Anxiety and Depression Scale (HADS) to measure anxiety and depression tendencies ^92^.

At the end of the stress task, patients also rated its characteristics (novelty, difficulty, stressfulness, controllability, and predictability) on 5 different 100-point Likert scales, with 0 = not at all, 50 = moderately, 100 = extremely.

To ensure the SBST did not induce long term effects, the patients were phone called the day after and the week after their participation in the study and asked about their recovery. Based on 10 items from the PTSD Checklist Civilian for DSM-5 (PCL-C) ^22^ they were questioned about any consequence the task could have induced in their daily life, their subjective stress and nervousness level since the experiment, any disturbing memories, dreams, or thoughts related to the study. They were also asked if thinking back to the study made them feel overwhelmed, if it induced increased heartbeat, sweating, pain, or difficulty breathing, if they tended to avoid talking about the study, had any new concentration or sleep difficulty, more negative emotions, and finally, their current stress and pain level on a numerical rating scale from 0 (not at all) to 10 (most intense). Previous experience with social benefits were also investigated (either if they already faced a medical expert, had paper/email interactions with social benefits or were never involved in social benefits procedures) and are described in Table 1. These results will be described in detail in further work.

**Table 1:** Demographic characteristics of the study population. BPI: Brief pain inventory, HADS: Hospital anxiety and depression scale, PCS: Pain catastrophizing scale, PSS: Perceived stress scale.

| Patients characteristics (N = 40) |  |
| --- | --- |
| **Age:** years mean ± SD (min-max) | 42 ± 11.2 (22 - 63) |
| **Diagnosis N (%)** |  |
| Ehlers-Danlos syndrome | 25 (62.5%) |
| Hypermobility spectrum disorder | 15 (37.5%) |
| **Social benefits experience N (%)** |  |
| Faced a medical expert | 26 (65%) |
| Involved in procedures without facing a medical expert | 6 (15%) |
| No previous experience | 8 (20%) |
| **BMI (kg/m<sup>2</sup>)** |  |
| Mean ± SD (min-max) | 28.8 ± 8 (17 - 45.9) |
| **Activity involving movement (h/week)** |  |
| Mean ± SD (min-max) | 9.3 ± 8.3 (1 - 40) |
| **Smoking N (%)** |  |
| Yes: (mean ± SD) | 21 (52.5%): (6.2 packs year ± 11.2) |
| No | 19 (47.5%) |
| **Education level N (%)** |  |
| Compulsory education | 2 (5%) |
| Apprenticeship | 17 (42.5%) |
| Highschool | 2 (5%) |
| University | 16 (40%) |
| Others | 2 (5%) |
| **Civil status N (%)** |  |
| In a relationship or married | 20 (50%) |
| Single or divorced | 19 (47.5%) |
| Widow | 1 (2.5%) |
| **Questionnaires: mean ± SD (min-max)** |  |
| BPI-severity | 5.47 ± 1.49 (2.5 - 9) |
| BPI-interference | 5.51 ± 1.87 (1 - 8.57) |
| HADS-anxiety [% above cutoff ≥ 8] | 1.53 ± 3.36 (4 - 18) [77.5%] |
| HADS-depression [% above cutoff ≥ 8] | 8.88 ± 3.30 (2 - 15) [67.5%] |
| PCS [% above cutoff ≥ 30] | 25.43 ± 9.84 (6 - 42) [40%] |
| PSS [% of high PSS ≥ 27] | 22.40 ± 5.97 (8 - 35) [30%] |

## Statistical analyses

### Data analyses

Given that the SBST is a new stress paradigm, no prior data existed for an a priori power calculation. Therefore, a sample size of N = 40 was estimated based on previous studies using the TSST^29^. Statistical analyses were performed using python v3.8.10 and R v4.3 under RStudio v2021.09.1-372. Results were considered significant at p<0.05. One-way repeated-measures ANOVAs with Greenhouse-Geisser sphericity correction were used to evaluate time effect on subjective stress and pain ratings, HRV, EDA and salivary cortisol and α-amylase concentrations. If the time effect was significant, post-hoc pairwise t-tests were performed with a Bonferroni correction for multiple comparisons. All the results are reported corrected. To test the validity of the new stress task, the post-hoc analysis focused on the following comparisons: baseline vs stress task and stress task vs recovery. The entire set of post-hoc comparisons is presented in supplementary material.

Repeated measures correlations ^4^ were conducted to test associations between paired measures repeatedly assessed during the task, i.e. stress VAS, LF/HF ratio and salivary biomarkers concentrations. As cortisol is produced with a delay, its repeated measures correlations with stress VAS were done with a 10- min shift (for instance: stress VAS at baseline was tested with cortisol level at anticipation).

Two tailed paired t-tests were conducted to compare Positive and Negative Affect scores before and after the SBST completion.

Stress induction tasks responders are typically determined by cortisol increases. It can be defined as a cortisol increase from baseline to peak of ≥ 1.5 nmol/l ^57^, or the 30% highest (vs. non-responder: lowest) difference between baseline and highest post-stress sample ^43^. Given the very low cortisol concentrations in our population, cortisol stress reactivity was defined as the change between the baseline level and the higher of the post-stress samples (upper and lower 30% Δ saliva cortisol among the 36 participants with sufficient saliva volume for analysis: N= 11 participants low, and N= 11 high).

### Salivary samples

An automated mathematical correction was implemented to account for the inherent diurnal variation in cortisol and α-amylase It aimed to isolate stress task-induced peaks, independently of the individual underlying natural variation during the day and considering very low signal in the measure of cortisol (mean baseline concentration = 2.06 ± 1.36 nmol/L vs. mean afternoon levels in healthy controls above 7 nmol/L^42,50^).

Cortisol and α-amylase levels were modeled as a function of time centered around the grand mean ^61^, as depicted in Figure 2, a). To capture the natural decrease in α-amylase and cortisol concentrations, we expressed the distribution of the salivary biomarkers’ values using polynomial terms over the reported time range and applied a quadratic regression to account for the natural variations. Stress-induced peaks (anticipation and SBST timepoints) were excluded not to affect the regression parameterization, as shown in Figure 2, b). By subtracting the resulting regression curves from the actual cortisol and α-amylase curves (a – b = c), we corrected for the natural diurnal evolution of these concentrations. The corrected results, called Δ cortisol (Figure 2, c) or Δ α-amylase are considered as the increase related to the task.

**Figure 2:**
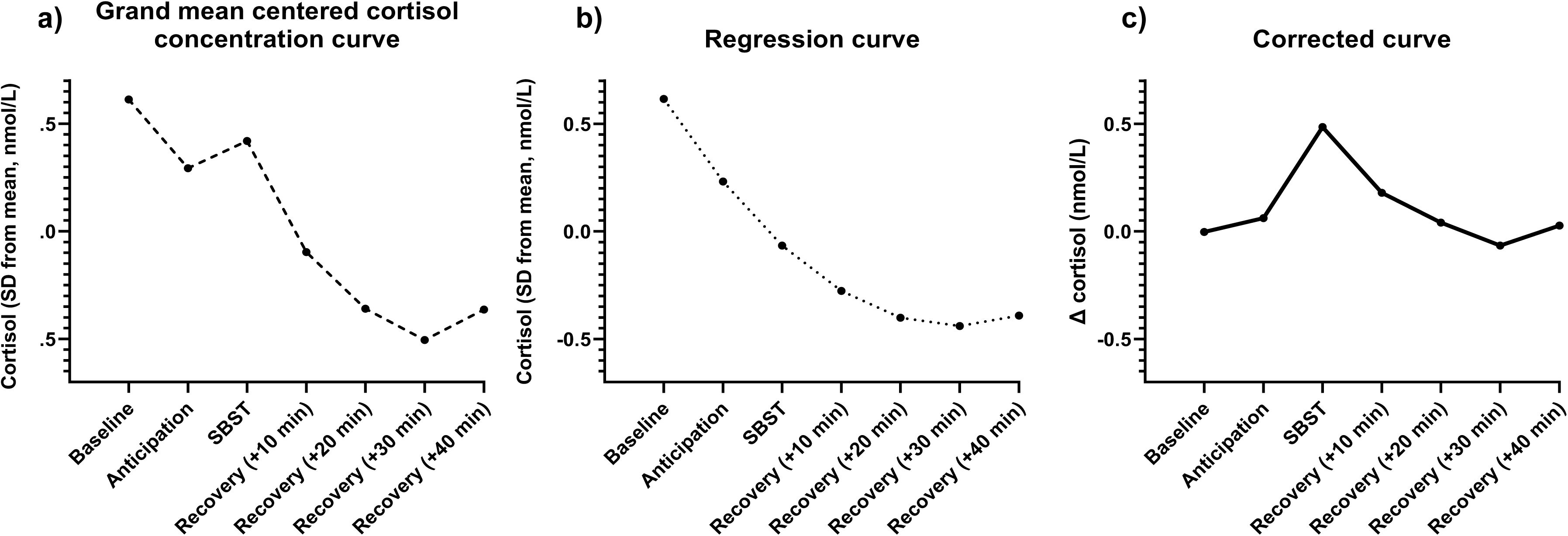
Example of the correction method for salivary cortisol and -amylase concentrations, here applied to data from one patient. a) The discontinued line represents a salivary cortisol concentration pattern that was fitted as a function of time centered around the grand mean. b) The dotted line represents a quadratic regression that was used to capture cortisol natural variations with an automatic peak removal. c) In bold, the resulting regression curve was subtracted from the actual cortisol curve to correct for the natural evolution of the concentration.

## Results

### Demographic data and psychological traits

The study sample consisted in 40 female subjects with an average age of 42 (±11.2 years). hEDS diagnostic criteria were fulfilled in 25, whereas 15 were categorized as suffering from HSD. Their demographic characteristics are described in Table 1.

On average, the group reported daily pain with a moderate intensity that considerably interfered with their daily life. A tendency to catastrophize about pain was also frequently observed. Anxiety and depression were likely co-morbidities, with 55% scoring above clinical detection criteria for both conditions. All patients but one displayed moderate to high subjective stress scores.

### Visual analog scales for subjective stress and subjective pain during the SBST

A significant time effect was reported for subjective stress (F(4, 155.98)=39.82, p<.0001, η²=.29) (see Figure 3, a). Compared to baseline (M=1.56, SD=1.72), subjective stress was significantly higher during the interview (M=4.84, SD=2.54; t(39)=8.69; p<.0001) and the arithmetic task (M=5.36 ; SD=2.69; p<.0001). Compared to the end of the arithmetic task, subjective stress was significantly lower at the end of the recovery phase (M=1.46; SD=1.77; p<.0001).

**Figure 3:**
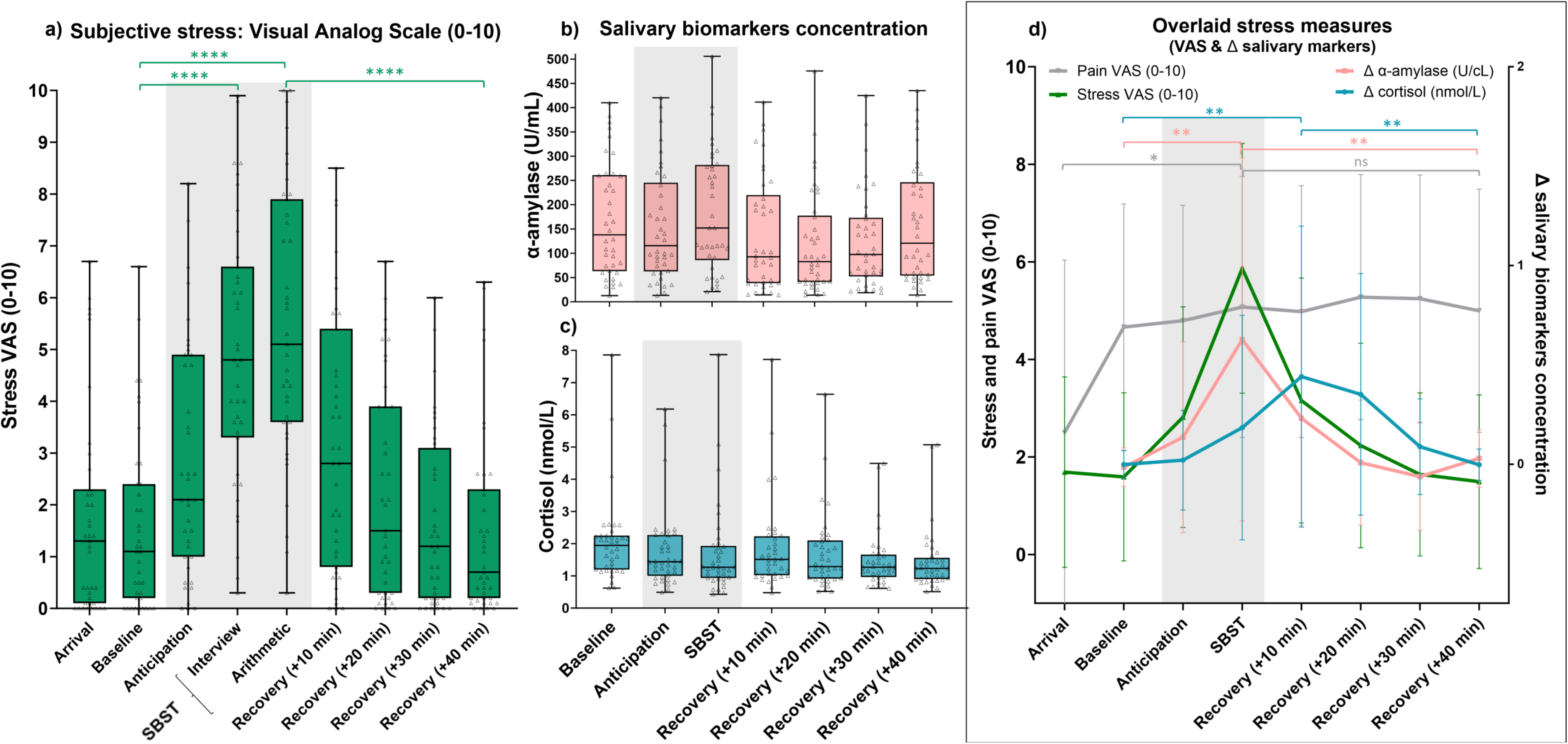
Representation of the measures of the stress response. a) Self-reported subjective stress (green, Visual Analog Scale -VAS), b) salivary α-amylase (U/mL, pink) and c) salivary cortisol (nmol/L, blue) concentrations measured during the different phases of the Social Benefits Stress Test (SBST) d) Stress response measures (VAS, Δ salivary cortisol (nmol/L) and Δ salivary α- amylase (U/cL)) displayed together during the SBST. As pre-determined, the presented statistics are the baseline vs stress-task and end-of-stress-task vs recovery post-hoc tests, the others can be found in supplementary. The whiskers represent the standard deviation, the grey frame show the timeframe where the task occurred, asterisks indicate significant reported post-hoc between successive SBST phases: **** for p < 0.0001, *** for p < 0.001, ** for p < 0.01 and * for p < 0.05

A significant time effect was also reported for the pain ratings (F(3.41, 133.04)=4.15, p=.005, η²=.013). Compared to arrival (M=4.32, SD=2.43), subjective pain was significantly higher during the arithmetic task (M=5.08, SD=2.68, p=.032). All the comparisons are presented in the supplementary material.

### Salivary α-amylase and cortisol

Two participants were excluded from all salivary analyses due to insufficient saliva volume (analysis for α- amylase n=38), and 2 more from the cortisol analyses because the concentrations were under detection levels (n=36). Raw salivary concentration are illustrated in figure 3, b) for the α-amylase and figure 3, c) for the cortisol.

During the task, a significant time effect was reported for the corrected Δ α-amylase concentration (F(2.7, 99.81)=1.36, p<.0001, η²=.183). Compared to baseline (M=1.52, SD=9.76), Δ α-amylase concentration was significantly higher during the stress task (M=62.76, SD=91.3; p=.003). Then, compared to the stress task, Δ α-amylase concentration was significantly lower after the recovery phase (M=2.96, SD=14.55; p=.006) (Figure 3, d).

A significant time effect was also reported for Δ salivary cortisol concentration (F(2.43, 85.15)=6.83, p<.0009, η²=.125). Compared to baseline (M=.002, SD=.07), Δ salivary cortisol concentration was significantly higher 10 min (M=.44, SD=.76; p<.026) and 20 min (M=.35, .61; p<.029) after the stress task. Δ salivary cortisol concentration was significantly lower at the end of the recovery phase (M=.003, SD=.08) compared to 20 min post-stress task (p<.026) Figure 3, d).

### Heart rate variability

Due to technical issues during the recordings, 6 participants with missing data were excluded from the these analyses (N=34). A significant time effect was reported for heart rate (F(2.52, 8.7)=26.99, p<.0001, η²=.12) (Figure 4, a). Compared to the baseline (M=78.34, SD=1.87), the heart rate was significantly higher during the interview (M=84.01, SD=13.3; p=.016) and arithmetic task (M=83.37, SD=12.36; p<.0006,). Heart rate was also significantly lower during the recovery phase (M=73.51, SD=1.1; p<.0001,) compared to arithmetic. We observed significant time effects for LF (F(4, 128)=9.4, p<.0001, η²=.184). Compared to the baseline (M=.016, SD=.01), LF was significantly higher during the arithmetic task (M=.031, SD=.014; p<.0001) and then decreased significantly at the end of the recovery phase (40 minutes) (M=.016, SD=.014; p=.003) (Figure 4,b).

**Figure 4:**
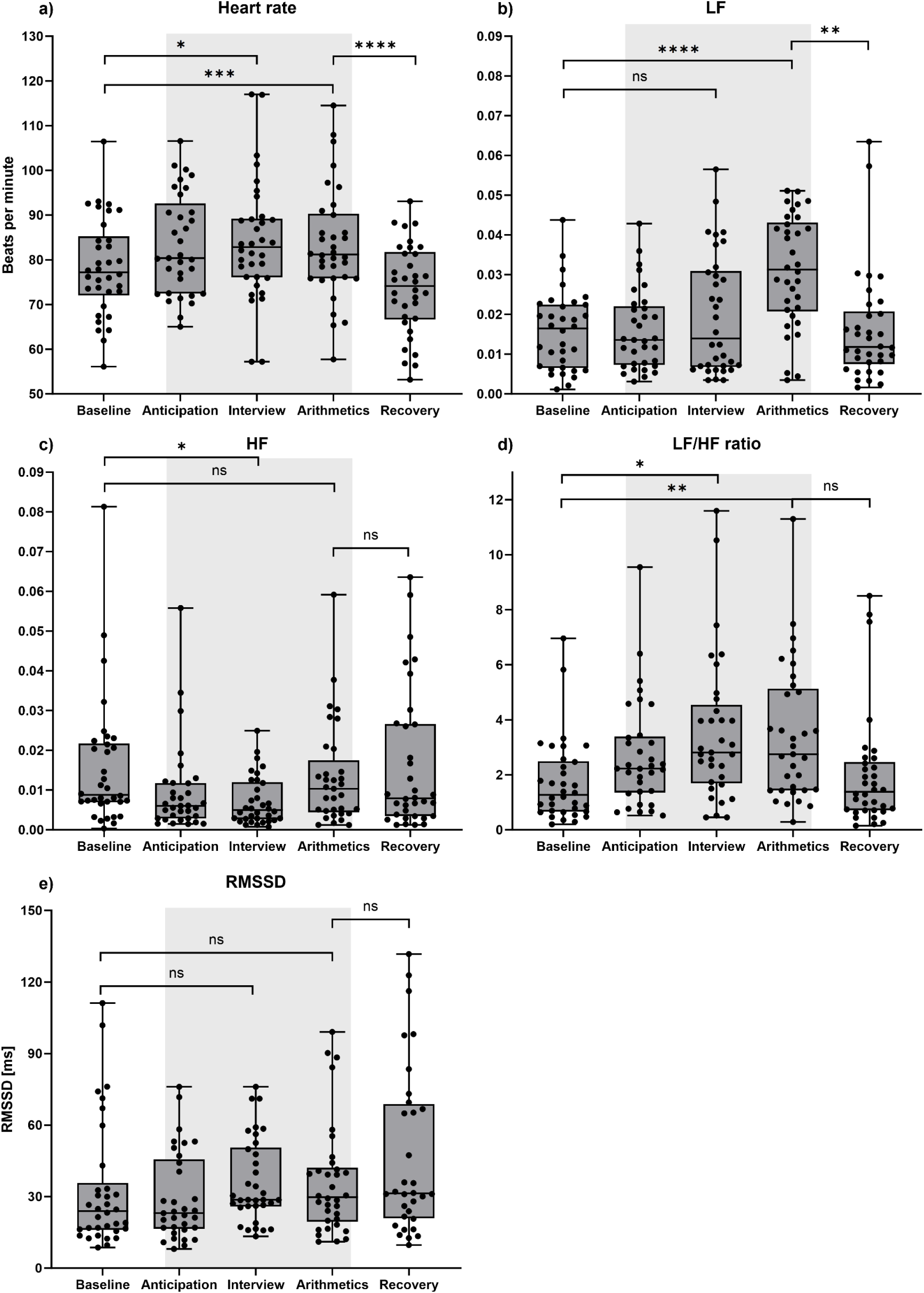
Heart rate variability metrics during the Social Benefits Stress Test. a) Heart rate (beats per minute) b) Low frequency (LF), c) High frequency (HF), d) LF/HF ratio (no unit) and e) Root Mean Square of Successive Differences (RMSSD, in ms) measured during the different phases of the Social Benefits Stress Test (SBST). As pre-determined, the presented statistics are the baseline vs stress-task and end-of-stress-task vs recovery post-hoc tests, the others can be found in supplementary. The whiskers represent the standard deviation, the grey area show the timeframe where the task occurred, asterisks indicate significant reported post-hoc between successive SBST phases: **** for p < 0.0001, *** for p < 0.001, ** for p< 0.01 and * for p < 0.05.

There was a significant time effect for HF (F(2.66, 79.74)=3.39, p=.027, η²=.08). HF was significantly lower during the interview (M=.007, SD=.006) compared to the baseline (M=.014, SD=.012; p=.045) (Figure 4,c).

A significant time effect was also observed for LF/HF ratio (F(2.66, 85.2)=7.07, p<.0005, η²=.13). LF/HF ratio was significantly higher during both the interview (M=3.98, SD=3.63; p=.015) and the arithmetic parts (M=3.82, SD=3.33; p=.007) compared to the baseline (M=1.74, SD=1.51, Figure 4, d).

For RMSSD, a significant time effect was observed (F(2.18, 69.78)=5.17, p<.007, η²=.083). However, there were no post-hoc differences after applying Bonferroni correction (Figure 4, e).

### Electrodermal activity

Due to technical issues during the recordings N=11 datasets were excluded from the these analyses, where the device did not record the EDA for 6 participants and delivered corrupted files/data with heavy artefacts in 5 (final sample N= 29). A significant time effect was detected for skin conductance levels (SCL) (F(1.89, 53.05) = 9.29, p < .001, η² = .04) (Figure 5, a). Compared to the baseline (M= 0.96, SD = 1.09), SCL was significantly higher during the interview (M = 2.34, SD = 2.76; p = .01) and arithmetic task (M = 2.1, SD = 2.67; p = .011). SCL was also significantly lower during the recovery phase (M = 1.60, SD = 2.05) compared to arithmetic (M = 2.09, SD = 2.67; p = .012).

**Figure 5:**
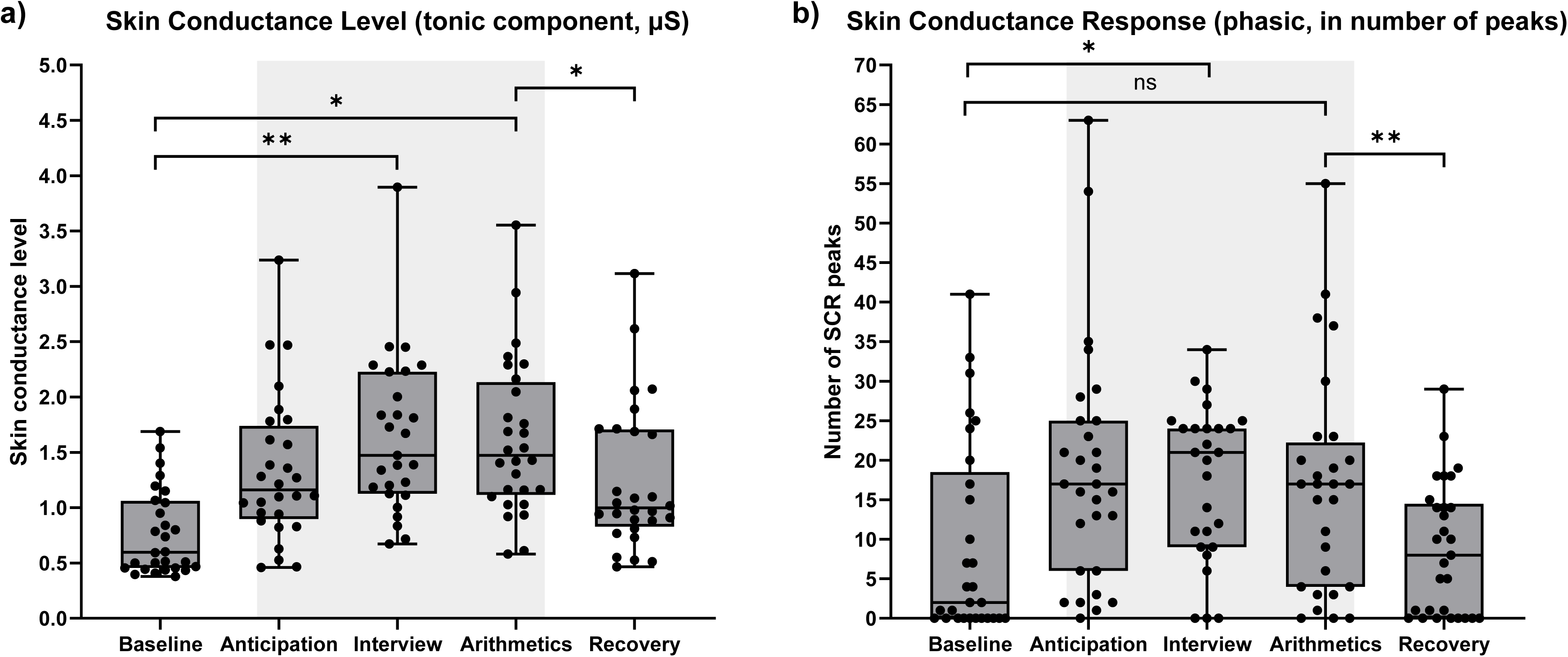
Electrodermal activity metrics during the Social Benefits Stress Test. a) Tonic component skin conductance level (in μS) b) Phasic component skin conductance response (in number of peaks) measured during the different phases of the Social Benefits Stress Test (SBST). The whiskers represent the standard deviation, the grey area show the timeframe where the task occurred, asterisks indicate significant reported post-hoc between successive SBST phases: **** for p < 0.0001, *** for p < 0.001, ** for p< 0.01 and * for p < 0.05. We present as stated in the method baseline-vs-stress-task and end-of-stress-task-vs-recovery post-hoc tests, others are in supplementary.

We observed significant time effects for skin conductance response (SCR) (F(3.06, 79.55) = 6.06, p < .001, η² = .12). Compared to the baseline (M = 9.31, SD = 12.32), the number of SCR peaks was significantly higher during the interview (M = 17.48, SD = 9.68; p = .049) and decreased significantly during the recovery phase (M = 8.76, SD = 8.34) compared to the arithmetic task (M = 16.54, SD = 13.88; p < .01) (Figure 5, b).

### Correlation between objective and subjective stress measures

Subjective stress VAS significantly correlated with: Δ salivary cortisol concentration (r_rm_(179)=.32, p<.0001), Δ α-amylase concentration (r_rm_(227)=.32, p<.0001), LF/HF ratio (r_rm_(97)=.28, p<.005) and the EDA tonic component SCL(r_rm_(115)=.27, p<.01). The EDA tonic component also correlated with LF/HF ratio (r_rm_(97) =.09, p < .05).

### Patients’ perception of the task, debriefing and post-task effects

At the end of the recovery, participants rated the task as difficult (M=79.4/100, SD=19.46), stressful (M=76.5, SD=21.64), moderately novel (M=50.4, SD=36.67), not so controllable (M=41.75, SD=29.32) and weakly predictable (M=20.88, SD=3.17).

There was a significant decrease in the PANAS Positive Affect after the task (M=24.08, SD=7.47 before the task vs M=21.63, SD=8.92 after; t(39)=-2.89, p=.006) and a significant increase in Negative Affects (M=13.06, SD=3.76 before vs. M=20.68, SD=7.90 after; t(39)=6.15, p<.001).

Finally, at the one-day follow-up call, participants reported what they considered as usual levels of stress (M=3.03/10, SD=2.53) and pain (M=5.65, SD=1.77) in their daily lives (n=38). Similar ratings were observed after one week (Stress: M=2.68, SD=2.01 and Pain: M=5.42, SD=2.62, n=34). Among the list of 16 potential aftereffects, participants reported only few symptoms (M=1.42 symptoms /16, SD=1.83 after one day and M=1.06, SD=2.01 after a week), The most common symptoms were fatigue/pain (n=15), nervosity (n=11) and thoughts regarding the task (n=5).

One participant experienced an exacerbation of a past trauma (evoked by the ocean documentary), but not specifically related to the stress task of the SBST. One other participant experienced sleep disturbance that she attributed to the task and to her having no more psychiatric support at the time, and for which she received counsel from the study physician.

### Exploratory outcomes

In exploratory analyses, we tested the effects of the following potential personal-level confounders on stress induction: age, education, diagnosis, pain (BPI severity and interference, maximum pain VAS, increase of pain VAS), psychological parameters (HADS, PCS), stress perception (PSS, BRS), work incapacity. Of these, none were significant, nor did it justify multilevel analyses. The statistics of these analyses are presented in the supplementary materials. Among the responders (n=11) and non-responders (n=11) based on Δ saliva cortisol, no psychological differences were found.

## Discussion

In this study, we developed and validated the Social Benefits Stress Test (SBST), an adaptation of the Trier Social Stress Test that is relevant to patients unable to work, for example due to chronic pain. The task design was informed by people with lived experience and the clinical experiences of diverse medical personnel treating people with chronic pain conditions. We tested if a simulated medical expert interview situation could be used as a relevant and ecological stressor in patients suffering from chronic widespread pain. We demonstrated that the SBST triggered a reversible moderate stress response, measured as significant and transient increases in self-reported stress (VAS), salivary markers (cortisol and α-amylase concentrations) and changes in heart rate and heart rate variability (HRV) and electrodermal activity (EDA) metrics.

### The SBST, a good stress induction task?

The SBST elicited a moderate to intense increase in subjective stress (mean +3.8 pts on VAS). This response was comparable or even higher than observed with the TSST in healthy volunteers, where a meta-analysis described a mean range of increase between 2.0 to 3.9 pts ^49^. In patients suffering from fibromyalgia, the TSST induced smaller increases in stress (mean increase: less than 1 point in n=27; and 2 points in, n=13) ^23,25^. Further studies using other stress paradigms in CWP patients (fibromyalgia), did not report the subjective stress increases (n=36, “Mental Arithmetic with Harassment) ^54^ or reported a smaller increase (n=23, +1.7 points for the two-choice reaction-time test) ^62^. The SBST also to lead to a significant increase in PANAS negative affect (NA) and decrease in positive affect (PA), both akin to the emotional response observed in healthy subjects during the TSST ^49^.

Moreover, the SBST successfully induced objective physiological responses. Heart rate temporarily increased during the stress task, akin to observations in fibromyalgia patients in TSST settings ^45^, and mental arithmetic exercises ^91^. The tonic and phasic EDA components were higher during the stress task, similarly to a prior observation of an SCL increase in fibromyalgia during an arithmetic task and social conflict^79^. We also observed a transient increase in Δ salivary cortisol reaching a peak between +10min and +20min after the stress induction, and a transient increase in Δ α-amylase during the task, mirroring the responses seen in individuals with rheumatoid arthritis, psoriasis or multiple sclerosis during the TSST ^3,26^. Importantly, we also observed that stress VAS positively correlated with the EDA tonic component and salivary biomarkers (Δ cortisol and Δ α-amylase). Such correlations between subjective and objective outcomes was not reported in chronic pain patients in most prior stress induction studies ^7,18,40^. Hence, this is supporting the strength of the SBST paradigm. In the TSST in other contexts, correlations between cortisol responses and subjective stress variables were found in 25% of the studies ^1,15^.

Furthermore, the SBST’s impact was transient, inducing an acute stress response without 1-week posttest effects except in two psychologically vulnerable patients (undisclosed PTSD n=1; lack of necessary psychological support n=1). Hence, this might require further caution in the eligibility criteria for further use of the SBST.

Previous studies have also adapted the TSST to choose an ecological stressor, more relevant and specific for a targeted population: recognizing their crying baby for new mothers, expressing how good a friend they are for adolescents or defending themselves against shoplifting accusations in adult men ^39,70,88^. In our study, the SBST demonstrates relevance for a chronic widespread pain population by considering their possible work impairment ^8^. Our manipulation check confirmed the task’s effectiveness in eliciting a challenging experience, which they rated as difficult, stressful, unpredictable, and uncontrollable. Yet, the novelty was rated as moderate, possibly indicating that the SBST’s realism resonated with participants who had undergone or heard about similar real medical expertise, or other medical encounters where they felt disbelief or judgment ^30^.

Finally, relying only on one confederate, the SBST presents a logistical advantage, compared to the panel of three confederates used in the original TSST.

### The stress response in chronic widespread pain

In our study and compared to prior published samples, hEDS/HSD patients exhibited very low salivary cortisol concentrations (i.e. M=2.06±1.36 nmol/L, compared to normative afternoon levels in healthy controls above 7 nmol/L ^42,50^), and elevated levels of α-amylase (i.e. M=168.54±117.19 U/mL, compared to mean afternoon levels under 140U/mL ^38^), suggesting an alteration of the neuroendocrine stress response system. To our knowledge there is little research describing the stress response in the hEDS/HSD population. The only identified publication on cortisol production in hEDS/HSD patients employed a small sample (n=10) without a unified collection time (9h-17h), hence not allowing a comparison with our data ^74^. We also observed a low HRV, that could be interpreted as reduced vagal activity or sympathetic hyperreactivity ^67^, congruent with the existing literature describing higher LF/HF ratio in a sample of hEDS/HSD (n=39) compared to controls ^28^. This is compatible with an autonomic dysfunction in hEDS/HSD ^53,55^. Further work with appropriate controls and the combination of biological and physiological markers will be needed to better characterize the specificities of the stress response in hEDS/HSD.

Baseline alterations in the stress response have been better studied in other patient populations with chronic widespread pain. In fibromyalgia, reduced cortisol awakening response ^68^, elevated levels of salivary α- amylase ^24,33^, altered autonomic regulation of electrodermal activity ^63^, and sympathetic mediated HRV at baseline have been described ^67,75^. In the light of these baseline alterations, the stress response induction by a stress task is of specific interest. Individuals with fibromyalgia compared to healthy controls exhibited a blunted cortisol response and higher subjective stress levels when exposed to the TSST ^23^. In our data, in the absence of a control population, whether of healthy volunteers or patients with another painful condition, it is not possible to qualify the observed stress response. Further investigations will be needed, including perhaps patients with different types of pain. This future research could be facilitated by the task validated here. Furthermore, a longer-term objective for patients suffering from chronic pain could be to support stress management and resilience promotion ^51^.

### Limitations

The SBST was created for the specific Swiss social benefits system, where medical experts are very frequently involved in deciding on the attribution of financial support. The connection between the simulated and real experiences will be described in further work. Nevertheless, the task could be easily adapted to match other countries’ social and or insurance systems, given the universal economic concerns and needs for medical, and social recognition as well as financial support amongst individuals struggling with disability due to chronic pain ^46^.

The study included only women, based on the heavily skewed hEDS/HSD demographics ^16^. It is of interest that in a prior stress induction task, women displayed a greater increase in subjective distress compared to men, but a lower increase in cortisol ^34,44^. Moreover, the pain experience is more frequently disbelieved in women than in men ^19^. Further work is needed to validate the SBST in men and examine potential sex-related difference in responses.

Pain ratings were higher after the SBST compared to arrival. However, the task design doesn’t allow to disentangle between a potential effect on pain of stress, vs. of immobilization and prolonged sitting. This would require a control task with similar patients seated without a stress induction.

Measurements of α amylase and cortisol may be accompanied by procedural limitations ^69^: for example low blood glucose levels and smoking are associated with blunted cortisol responses and lower α-amylase activity ^76^. We limited the effects of these factors by asking patients to have an early lunch (before noon) since the study started at 2pm, and not to drink anything other than water, nor smoke before the start of the experiment.

Since the experiment took place in the early afternoon, the very low cortisol levels and the diurnal decrease made the interpretation of the raw data difficult. Hence, we applied a correction method based on the subtraction of the estimated natural slope to isolate the task-induced peak ^57^. Instead of this estimation, a measure of the individual natural variations in diurnal cortisol and α-amylase, collected on another day (e.g. the day before the task) could be another option for the correction. This is to be considered for future studies.

## Conclusion

The present study developed the SBST, a new social stress task, and validated it in a chronic widespread pain population of individuals with hypermobility conditions. This lays the groundwork for future in depth investigations in chronic pain populations into stress responses, and in parallel, stress management and resilience. The SBST could also serve to test therapeutic tools targeting a potential abnormal stress response. Finally, this test could be used to quantify the stress involved in medical expertise processes and attract attention towards this still neglected issue.

## Acknowledgments

We thank Emma de Francesco and Mélanie Colin for their precious help in physiological data cleaning, Jérôme Savary for the development of the automated mathematic correction method for salivary biomarkers, Pr. Antje Horsch for her helpful insights in the task design and preliminary data analysis, and Dr. Vania Sandoz for sharing tools which helped in the practical implementation of the SBST.

