## Supplementary material for "Justify your disability! A simulated medical evaluation as a robust novel stress induction paradigm in chronic pain patients"

| VAS stress | | | | | | | | |
|---|---|---|---|---|---|---|---|---|
| **group1** | **group2** | **n1** | **n2** | **statistic** | **df** | **p** | **p.adj** | **p.adj.signif** |
| Anticipation | Arithmetics | 40 | 40 | -7,231637486 | 39 | 1,03E-08 | 0,000000371 | **** |
| Anticipation | Arrival | 40 | 40 | 3,897651498 | 39 | 0,000371 | 0,013 | * |
| Anticipation | Baseline | 40 | 40 | 5,214560763 | 39 | 0,00000635 | 0,000229 | *** |
| Anticipation | Interview | 40 | 40 | -7,182055022 | 39 | 1,21E-08 | 0,000000436 | **** |
| Anticipation | Recovery_10 | 40 | 40 | -0,994522651 | 39 | 0,326 | 1 | ns |
| Anticipation | Recovery_20 | 40 | 40 | 1,712295469 | 39 | 0,095 | 1 | ns |
| Anticipation | Recovery_30 | 40 | 40 | 3,608280686 | 39 | 0,000866 | 0,031 | * |
| Anticipation | Recovery_40 | 40 | 40 | 4,188499036 | 39 | 0,000155 | 0,006 | ** |
| Arithmetics | Arrival | 40 | 40 | 8,455233549 | 39 | 2,36E-10 | 8,5E-09 | **** |
| **Arithmetics** | **Baseline** | **40** | **40** | **8,959127057** | **39** | **5,23E-11** | **1,88E-09** | **** |
| Arithmetics | Interview | 40 | 40 | 2,014232428 | 39 | 0,051 | 1 | ns |
| Arithmetics | Recovery_10 | 40 | 40 | 6,373198477 | 39 | 0,000000157 | 0,00000565 | **** |
| Arithmetics | Recovery_20 | 40 | 40 | 7,542518851 | 39 | 3,9E-09 | 0,00000014 | **** |
| Arithmetics | Recovery_30 | 40 | 40 | 9,261241913 | 39 | 2,15E-11 | 7,74E-10 | **** |
| **Arithmetics** | **Recovery_40** | **40** | **40** | **9,433403768** | **39** | **1,3E-11** | **4,68E-10** | **** |
| Arrival | Baseline | 40 | 40 | 0,816435721 | 39 | 0,419 | 1 | ns |
| Arrival | Interview | 40 | 40 | -7,905709905 | 39 | 1,27E-09 | 4,57E-08 | **** |
| Arrival | Recovery_10 | 40 | 40 | -4,032059382 | 39 | 0,000249 | 0,009 | ** |
| Arrival | Recovery_20 | 40 | 40 | -1,700184665 | 39 | 0,097 | 1 | ns |
| Arrival | Recovery_30 | 40 | 40 | 0,327552659 | 39 | 0,745 | 1 | ns |
| Arrival | Recovery_40 | 40 | 40 | 0,804442387 | 39 | 0,426 | 1 | ns |
| **Baseline** | **Interview** | **40** | **40** | **-8,692291134** | **39** | **1,16E-10** | **4,18E-09** | **** |
| Baseline | Recovery_10 | 40 | 40 | -4,527241803 | 39 | 0,000055 | 0,002 | ** |
| Baseline | Recovery_20 | 40 | 40 | -2,25339544 | 39 | 0,03 | 1 | ns |
| Baseline | Recovery_30 | 40 | 40 | -0,209912142 | 39 | 0,835 | 1 | ns |
| Baseline | Recovery_40 | 40 | 40 | 0,390907006 | 39 | 0,698 | 1 | ns |
| Interview | Recovery_10 | 40 | 40 | 5,413751027 | 39 | 0,00000337 | 0,000121 | *** |
| Interview | Recovery_20 | 40 | 40 | 7,151688741 | 39 | 1,33E-08 | 0,000000479 | **** |
| Interview | Recovery_30 | 40 | 40 | 8,912731006 | 39 | 6E-11 | 2,16E-09 | **** |
| Interview | Recovery_40 | 40 | 40 | 8,415544912 | 39 | 2,66E-10 | 9,58E-09 | **** |
| Recovery_10 | Recovery_20 | 40 | 40 | 3,03832908 | 39 | 0,004 | 0,152 | ns |
| Recovery_10 | Recovery_30 | 40 | 40 | 4,710867354 | 39 | 0,000031 | 0,001 | ** |
| Recovery_10 | Recovery_40 | 40 | 40 | 3,944440063 | 39 | 0,000323 | 0,012 | * |
| Recovery_20 | Recovery_30 | 40 | 40 | 3,982797856 | 39 | 0,000288 | 0,01 | * |
| Recovery_20 | Recovery_40 | 40 | 40 | 2,484528492 | 39 | 0,017 | 0,626 | ns |
| Recovery_30 | Recovery_40 | 40 | 40 | 0,628067792 | 39 | 0,534 | 1 | ns |

| VAS pain | | | | | | | | |
|---|---|---|---|---|---|---|---|---|
| group1 | group2 | n1 | n2 | statistic | df | p | p.adj | p.adj.signif |
| Anticipation | Arithmetics | 40 | 40 | -1,377191539 | 39 | 0,176 | 1 | ns |
| Anticipation | Arrival | 40 | 40 | 2,763497773 | 39 | 0,009 | 0,312 | ns |
| Anticipation | Baseline | 40 | 40 | 1,222428879 | 39 | 0,229 | 1 | ns |
| Anticipation | Interview | 40 | 40 | -0,953903872 | 39 | 0,346 | 1 | ns |
| Anticipation | Recovery_10 | 40 | 40 | -0,89234213 | 39 | 0,378 | 1 | ns |
| Anticipation | Recovery_20 | 40 | 40 | -2,112808582 | 39 | 0,041 | 1 | ns |
| Anticipation | Recovery_30 | 40 | 40 | -1,877467134 | 39 | 0,068 | 1 | ns |
| Anticipation | Recovery_40 | 40 | 40 | -0,849986893 | 39 | 0,401 | 1 | ns |
| Arithmetics | Arrival | 40 | 40 | 3,602330409 | 39 | 0,000881 | 0,032 | * |
| **Arithmetics** | **Baseline** | **40** | **40** | **1,889828722** | **39** | **0,066** | **1** | **ns** |
| Arithmetics | Interview | 40 | 40 | 1,012093965 | 39 | 0,318 | 1 | ns |
| Arithmetics | Recovery_10 | 40 | 40 | 0,531390912 | 39 | 0,598 | 1 | ns |
| Arithmetics | Recovery_20 | 40 | 40 | -0,958908361 | 39 | 0,344 | 1 | ns |
| Arithmetics | Recovery_30 | 40 | 40 | -0,68635085 | 39 | 0,497 | 1 | ns |
| **Arithmetics** | **Recovery_40** | **40** | **40** | **0,335469815** | **39** | **0,739** | **1** | **ns** |
| Arrival | Baseline | 40 | 40 | -1,78444819 | 39 | 0,082 | 1 | ns |
| Arrival | Interview | 40 | 40 | -3,079096449 | 39 | 0,004 | 0,136 | ns |
| Arrival | Recovery_10 | 40 | 40 | -2,854411615 | 39 | 0,007 | 0,247 | ns |
| Arrival | Recovery_20 | 40 | 40 | -3,490563308 | 39 | 0,001 | 0,044 | * |
| Arrival | Recovery_30 | 40 | 40 | -3,128155336 | 39 | 0,003 | 0,12 | ns |
| Arrival | Recovery_40 | 40 | 40 | -2,405034989 | 39 | 0,021 | 0,756 | ns |
| **Baseline** | **Interview** | **40** | **40** | **-1,618443819** | **39** | **0,114** | **1** | **ns** |
| Baseline | Recovery_10 | 40 | 40 | -1,563410752 | 39 | 0,126 | 1 | ns |
| Baseline | Recovery_20 | 40 | 40 | -2,725476237 | 39 | 0,01 | 0,345 | ns |
| Baseline | Recovery_30 | 40 | 40 | -2,506119424 | 39 | 0,016 | 0,594 | ns |
| Baseline | Recovery_40 | 40 | 40 | -1,442348789 | 39 | 0,157 | 1 | ns |
| Interview | Recovery_10 | 40 | 40 | -0,299284116 | 39 | 0,766 | 1 | ns |
| Interview | Recovery_20 | 40 | 40 | -1,607340785 | 39 | 0,116 | 1 | ns |
| Interview | Recovery_30 | 40 | 40 | -1,344976637 | 39 | 0,186 | 1 | ns |
| Interview | Recovery_40 | 40 | 40 | -0,301117675 | 39 | 0,765 | 1 | ns |
| Recovery_10 | Recovery_20 | 40 | 40 | -2,210761867 | 39 | 0,033 | 1 | ns |
| Recovery_10 | Recovery_30 | 40 | 40 | -1,411696553 | 39 | 0,166 | 1 | ns |
| Recovery_10 | Recovery_40 | 40 | 40 | -0,079466763 | 39 | 0,937 | 1 | ns |
| Recovery_20 | Recovery_30 | 40 | 40 | 0,292891935 | 39 | 0,771 | 1 | ns |
| Recovery_20 | Recovery_40 | 40 | 40 | 1,79312756 | 39 | 0,081 | 1 | ns |
| Recovery_30 | Recovery_40 | 40 | 40 | 2,7126945 | 39 | 0,01 | 0,356 | ns |

| Δ cortisol | | | | | | | | |
|---|---|---|---|---|---|---|---|---|
| group1 | group2 | n1 | n2 | statistic | df | p | p.adj | p.adj.signif |
| Anticipation | Baseline | 36 | 36 | 0,452719776 | 35 | 0,654 | 1 | ns |
| Anticipation | Recovery_40 | 36 | 36 | 0,625678511 | 35 | 0,536 | 1 | ns |
| Anticipation | SBST | 36 | 36 | -1,473403822 | 35 | 0,15 | 1 | ns |
| Anticipation | Recovery_10 | 36 | 36 | -3,115852448 | 35 | 0,004 | 0,077 | ns |
| Anticipation | Recovery_30 | 36 | 36 | -1,018568338 | 35 | 0,315 | 1 | ns |
| Anticipation | Recovery_20 | 36 | 36 | -2,993349003 | 35 | 0,005 | 0,106 | ns |
| Baseline | Recovery_40 | 36 | 36 | 0,055680416 | 35 | 0,956 | 1 | ns |
| Baseline | SBST | 36 | 36 | -2,048305646 | 35 | 0,048 | 1 | ns |
| **Baseline** | **Recovery_10** | **36** | **36** | **-3,514472227** | **35** | **0,001** | **0,026** | * |
| Baseline | Recovery_30 | 36 | 36 | -2,392729561 | 35 | 0,022 | 0,466 | ns |
| Baseline | Recovery_20 | 36 | 36 | -3,474698848 | 35 | 0,001 | 0,029 | * |
| Recovery_40 | SBST | 36 | 36 | -1,953559224 | 35 | 0,059 | 1 | ns |
| **Recovery_40** | **Recovery_10** | **36** | **36** | **-3,555386831** | **35** | **0,001** | **0,023** | * |
| Recovery_40 | Recovery_30 | 36 | 36 | -1,849907314 | 35 | 0,073 | 1 | ns |
| Recovery_40 | Recovery_20 | 36 | 36 | -3,517808627 | 35 | 0,001 | 0,026 | * |
| SBST | Recovery_10 | 36 | 36 | -1,986179915 | 35 | 0,055 | 1 | ns |
| SBST | Recovery_30 | 36 | 36 | 0,890004759 | 35 | 0,38 | 1 | ns |
| SBST | Recovery_20 | 36 | 36 | -1,354844847 | 35 | 0,184 | 1 | ns |
| Recovery_10 | Recovery_30 | 36 | 36 | 2,731926829 | 35 | 0,01 | 0,206 | ns |
| Recovery_10 | Recovery_20 | 36 | 36 | 1,38052416 | 35 | 0,176 | 1 | ns |
| Recovery_30 | Recovery_20 | 36 | 36 | -2,785280174 | 35 | 0,009 | 0,18 | ns |

| Δ α-amylase | | | | | | | | |
|---|---|---|---|---|---|---|---|---|
| group1 | group2 | n1 | n2 | statistic | df | p | p.adj | p.adj.signif |
| Anticipation | Baseline | 38 | 38 | 1,817487109 | 37 | 0,077 | 1 | ns |
| Anticipation | Recovery_40 | 38 | 38 | 1,328247541 | 37 | 0,192 | 1 | ns |
| Anticipation | SBST | 38 | 38 | -3,458254565 | 37 | 0,001 | 0,029 | * |
| Anticipation | Recovery_10 | 38 | 38 | -0,947812303 | 37 | 0,349 | 1 | ns |
| Anticipation | Recovery_30 | 38 | 38 | 2,213307178 | 37 | 0,033 | 0,695 | ns |
| Anticipation | Recovery_20 | 38 | 38 | 1,279767602 | 37 | 0,209 | 1 | ns |
| Baseline | Recovery_40 | 38 | 38 | -1,205366465 | 37 | 0,236 | 1 | ns |
| **Baseline** | **SBST** | **38** | **38** | **-4,257271185** | **37** | **0,000136** | **0,003** | ** |
| Baseline | Recovery_10 | 38 | 38 | -2,595996603 | 37 | 0,013 | 0,281 | ns |
| Baseline | Recovery_30 | 38 | 38 | 1,303714587 | 37 | 0,2 | 1 | ns |
| Baseline | Recovery_20 | 38 | 38 | -0,49919264 | 37 | 0,621 | 1 | ns |
| **Recovery_40** | **SBST** | **38** | **38** | **-4,036734651** | **37** | **0,000262** | **0,006** | ** |
| Recovery_40 | Recovery_10 | 38 | 38 | -2,486807379 | 37 | 0,018 | 0,368 | ns |
| Recovery_40 | Recovery_30 | 38 | 38 | 1,382767887 | 37 | 0,175 | 1 | ns |
| Recovery_40 | Recovery_20 | 38 | 38 | 0,368258282 | 37 | 0,715 | 1 | ns |
| SBST | Recovery_10 | 38 | 38 | 2,9540989 | 37 | 0,005 | 0,114 | ns |
| SBST | Recovery_30 | 38 | 38 | 4,424244761 | 37 | 0,0000822 | 0,002 | ** |
| SBST | Recovery_20 | 38 | 38 | 3,920288192 | 37 | 0,000368 | 0,008 | ** |
| Recovery_10 | Recovery_30 | 38 | 38 | 2,663901013 | 37 | 0,011 | 0,239 | ns |
| Recovery_10 | Recovery_20 | 38 | 38 | 2,129629009 | 37 | 0,04 | 0,838 | ns |
| Recovery_30 | Recovery_20 | 38 | 38 | -1,00491284 | 37 | 0,321 | 1 | ns |

#### EDA number of peaks

| group1 | group2 | n1 | n2 | statistic | df | p | p.adj | p.adj.signif |
|---|---|---|---|---|---|---|---|---|
| Anticipation | Arithmetics | 29 | 29 | 0,647459 | 27 | 0,523 | 1 | ns |
| Anticipation | Baseline | 29 | 29 | 2,962547 | 28 | 0,006 | 0,062 | ns |
| Anticipation | Interview | 29 | 29 | 0,072713 | 26 | 0,943 | 1 | ns |
| Anticipation | Recovery | 29 | 29 | 3,524582 | 28 | 0,001 | 0,015 | * |
| **Arithmetics** | **Baseline** | **29** | **29** | **2,431762** | **27** | **0,022** | **0,219** | **ns** |
| Arithmetics | Interview | 29 | 29 | -1,04009 | 26 | 0,308 | 1 | ns |
| **Arithmetics** | **Recovery** | **29** | **29** | **3,95496** | **27** | **0,000499** | **0,005** | ****** |
| **Baseline** | **Interview** | **29** | **29** | **-3,07514** | **26** | **0,005** | **0,049** | ***** |
| Baseline | Recovery | 29 | 29 | 0,195317 | 28 | 0,847 | 1 | ns |
| Interview | Recovery | 29 | 29 | 5,053711 | 26 | 2,92E-05 | 0,00029 | *** |

#### EDA SCL

| group1 | group2 | n1 | n2 | statistic | df | p | p.adj | p.adj.signif |
|---|---|---|---|---|---|---|---|---|
| Anticipation | Arithmetics | 29 | 29 | -3,49296 | 28 | 0,002 | 0,016 | * |
| Anticipation | Baseline | 29 | 29 | 2,675099 | 28 | 0,012 | 0,123 | ns |
| Anticipation | Interview | 29 | 29 | -2,01018 | 28 | 0,054 | 0,541 | ns |
| Anticipation | Recovery | 29 | 29 | 1,541425 | 28 | 0,134 | 1 | ns |
| **Arithmetics** | **Baseline** | **29** | **29** | **3,645696** | **28** | **0,001** | **0,011** | ***** |
| Arithmetics | Interview | 29 | 29 | -1,03194 | 28 | 0,311 | 1 | ns |
| **Arithmetics** | **Recovery** | **29** | **29** | **3,620448** | **28** | **0,001** | **0,012** | ***** |
| **Baseline** | **Interview** | **29** | **29** | **-3,68154** | **28** | **0,00098** | **0,01** | ****** |
| Baseline | Recovery | 29 | 29 | -3,24084 | 28 | 0,003 | 0,031 | * |
| Interview | Recovery | 29 | 29 | 3,0931 | 28 | 0,004 | 0,045 | * |

| Heart rate | | | | | | | | |
|---|---|---|---|---|---|---|---|---|
| **group1** | **group2** | **n1** | **n2** | **statistic** | **df** | **p** | **p.adj** | **p.adj.signif** |
| Anticipation | Arithmetics | 34 | 34 | -0,760182299 | 32 | 0,453 | 1 | ns |
| Anticipation | Baseline | 34 | 34 | 5,539636523 | 32 | 0,00000413 | 0,0000413 | **** |
| Anticipation | Interview | 34 | 34 | -1,162889512 | 32 | 0,253 | 1 | ns |
| Anticipation | Recovery | 34 | 34 | 10,38070076 | 32 | 9,02E-12 | 9,02E-11 | **** |
| **Arithmetics** | **Baseline** | **34** | **34** | **4,61671773** | **33** | **0,0000568** | **0,000568** | ******* |
| Arithmetics | Interview | 34 | 34 | -0,493945409 | 33 | 0,625 | 1 | ns |
| **Arithmetics** | **Recovery** | **34** | **34** | **7,566759181** | **33** | **1,05E-08** | **0,000000105** | ******** |
| **Baseline** | **Interview** | **34** | **34** | **-3,432682852** | **33** | **0,002** | **0,016** | ***** |
| Baseline | Recovery | 34 | 34 | 5,551139745 | 33 | 0,00000362 | 0,0000362 | **** |
| Interview | Recovery | 34 | 34 | 7,001899638 | 33 | 5,23E-08 | 0,000000523 | **** |

| LF | | | | | | | | |
|---|---|---|---|---|---|---|---|---|
| **group1** | **group2** | **n1** | **n2** | **statistic** | **df** | **p** | **p.adj** | **p.adj.signif** |
| Anticipation | Arithmetics | 34 | 34 | -5,280552893 | 32 | 0,00000878 | 0,0000878 | **** |
| Anticipation | Baseline | 34 | 34 | 0,225571359 | 32 | 0,823 | 1 | ns |
| Anticipation | Interview | 34 | 34 | -1,381953077 | 32 | 0,177 | 1 | ns |
| Anticipation | Recovery | 34 | 34 | 0,087392289 | 32 | 0,931 | 1 | ns |
| **Arithmetics** | **Baseline** | **34** | **34** | **5,417322024** | **33** | **0,00000538** | **0,0000538** | ******** |
| Arithmetics | Interview | 34 | 34 | 4,690185928 | 33 | 0,0000458 | 0,000458 | *** |
| **Arithmetics** | **Recovery** | **34** | **34** | **4,059779114** | **33** | **0,000284** | **0,003** | ****** |
| **Baseline** | **Interview** | **34** | **34** | **-1,372216046** | **33** | **0,179** | **1** | **ns** |
| Baseline | Recovery | 34 | 34 | -0,123030482 | 33 | 0,903 | 1 | ns |
| Interview | Recovery | 34 | 34 | 0,972544566 | 33 | 0,338 | 1 | ns |

| HF | | | | | | | | |
|---|---|---|---|---|---|---|---|---|
| **group1** | **group2** | **n1** | **n2** | **statistic** | **df** | **p** | **p.adj** | **p.adj.signif** |
| Anticipation | Arithmetics | 34 | 34 | -1,783184394 | 32 | 0,085 | 0,847 | ns |
| Anticipation | Baseline | 34 | 34 | -1,894826054 | 32 | 0,068 | 0,678 | ns |
| Anticipation | Interview | 34 | 34 | 0,742122055 | 32 | 0,464 | 1 | ns |
| Anticipation | Recovery | 34 | 34 | -2,480255899 | 32 | 0,019 | 0,19 | ns |
| **Arithmetics** | **Baseline** | **34** | **34** | **-0,300621439** | **33** | **0,766** | **1** | **ns** |
| Arithmetics | Interview | 34 | 34 | 2,648420049 | 33 | 0,013 | 0,126 | ns |
| **Arithmetics** | **Recovery** | **34** | **34** | **-0,780570511** | **33** | **0,441** | **1** | **ns** |
| **Baseline** | **Interview** | **34** | **34** | **3,063976876** | **33** | **0,004** | **0,045** | ***** |
| Baseline | Recovery | 34 | 34 | -0,565228073 | 33 | 0,576 | 1 | ns |
| Interview | Recovery | 34 | 34 | -2,62022862 | 33 | 0,014 | 0,135 | ns |

| LF/HF ratio | | | | | | | | |
|---|---|---|---|---|---|---|---|---|
| **group1** | **group2** | **n1** | **n2** | **statistic** | **df** | **p** | **p.adj** | **p.adj.signif** |
| Anticipation | Arithmetics | 34 | 34 | -1,825618363 | 32 | 0,077 | 0,773 | ns |
| Anticipation | Baseline | 34 | 34 | 2,690423819 | 32 | 0,011 | 0,112 | ns |
| Anticipation | Interview | 34 | 34 | -2,123411937 | 32 | 0,042 | 0,415 | ns |
| Anticipation | Recovery | 34 | 34 | 2,415194839 | 32 | 0,022 | 0,216 | ns |
| **Arithmetics** | **Baseline** | **34** | **34** | **3,761172195** | **33** | **0,000659** | **0,007** | ****** |
| Arithmetics | Interview | 34 | 34 | -0,21599152 | 33 | 0,83 | 1 | ns |
| **Arithmetics** | **Recovery** | **34** | **34** | **2,511886547** | **33** | **0,017** | **0,171** | **ns** |
| **Baseline** | **Interview** | **34** | **34** | **-3,472339689** | **33** | **0,001** | **0,015** | ***** |
| Baseline | Recovery | 34 | 34 | -0,751898395 | 33 | 0,457 | 1 | ns |
| Interview | Recovery | 34 | 34 | 2,820148649 | 33 | 0,008 | 0,081 | ns |

| RMSSD | | | | | | | | |
|---|---|---|---|---|---|---|---|---|
| **group1** | **group2** | **n1** | **n2** | **statistic** | **df** | **p** | **p.adj** | **p.adj.signif** |
| Anticipation | Arithmetics | 34 | 34 | -1,356168331 | 32 | 0,185 | 1 | ns |
| Anticipation | Baseline | 34 | 34 | -0,414159632 | 32 | 0,682 | 1 | ns |
| Anticipation | Interview | 34 | 34 | -2,000829799 | 32 | 0,054 | 0,54 | ns |
| Anticipation | Recovery | 34 | 34 | -3,212410495 | 32 | 0,003 | 0,03 | * |
| **Arithmetics** | **Baseline** | **34** | **34** | **0,835059707** | **33** | **0,41** | **1** | **ns** |
| Arithmetics | Interview | 34 | 34 | 0,079706595 | 33 | 0,937 | 1 | ns |
| **Arithmetics** | **Recovery** | **34** | **34** | **-2,389959119** | **33** | **0,023** | **0,227** | **ns** |
| **Baseline** | **Interview** | **34** | **34** | **-0,689837693** | **33** | **0,495** | **1** | **ns** |
| Baseline | Recovery | 34 | 34 | -3,131273695 | 33 | 0,004 | 0,036 | * |
| Interview | Recovery | 34 | 34 | -2,560975172 | 33 | 0,015 | 0,152 | ns |

| **HADS Anxiety** | |
|---|---|
| One-way ANOVA | |
| F | 0,0343 |
| df1 | 2 |
| df2 | 20,5 |
| p | 0.966 |
| **HADS Depression** | |
| One-way ANOVA | |
| F | 2,63 |
| df1 | 2 |
| df2 | 22,4 |
| p | 0,094 |
| **Perceived Stress Scale** | |
| One-way ANOVA | |
| F | 0,605 |
| df1 | 2 |
| df2 | 8,9 |
| p | 0,567 |
| **Brief Resilience Scale** | |
| One-way ANOVA | |
| F | 0,195 |
| df1 | 2 |
| df2 | 5,54 |
| p | 0,828 |

| **BPI Pain Severity** | |
|---|---|
| Independent Samples t-test | |
| Statistic | -0,847 |
| df | 38 |
| p | 0,402 |
| **BPI Pain Interference** | |
| Independent Samples t-test | |
| Statistic | -1,83 |
| df | 38 |
| p | 0,075 |
| **Catastrophizing** | |
| Independent Samples t-test | |
| Statistic | -1,24 |
| df | 38 |
| p | 0,224 |
| **Diagnosis** | |
| Independent Samples t-test | |
| Statistic | -0,479 |
| df | 38 |
| p | 0,634 |
| **Age** | |
| Person correlation | |
| Pearson's r | -0,033 |
| p | 0,838 |

| **Education** | |
|---|---|
| Independent Samples t-test | |
| Statistic | -0,927 |
| df | 38 |
| p | 0,36 |
| **Maximum pain VAS** | |
| Independent Samples t-test | |
| Statistic | -0,983 |
| df | 38 |
| p | 0,332 |
| **Increase pain VAS** | |
| Independent Samples t-test | |
| Statistic | 0,273 |
| df | 38 |
| p | 0,786 |
| **Work incapacity** | |
| Independent Samples t-test | |
| Statistic | -0,734 |
| df | 38 |
| p | 0,467 |

| **Paired Samples T-Test between Responders (R) and Non-Responders (NR)** | | | | |
|---|---|---|---|---|
| Comparison | | statistic | df | p |
| R_BPI severity | NR_BPI severity | -0.0585 | 10.0 | 0.954 |
| R_BPI interference | NR_BPI interference | 1.0669 | 10.0 | 0.311 |
| R_Resilience | NR_Resilience | -1.2290 | 10.0 | 0.247 |
| R_HADS Anxiety | NR_HADS Anxiety | 1.5316 | 10.0 | 0.157 |
| R_HADS Depression | NR_HADS Depression | 1.7754 | 10.0 | 0.106 |
| R_Catastrophizing | NR_Catastrophizing | 0.5906 | 10.0 | 0.568 |
| R_PSS | NR_PSS | -0.0313 | 10.0 | 0.976 |

N° patient :

Experimenter 2 is seated, rising when the patient arrives. During the experiment, she takes notes on the participant's behavior, while adopting a skeptical attitude. She must express doubts and subtle guilt, and not appear to believe or be convinced by the relevance and veracity of the participant's words.
**Attitude is the key to SBST.**

### 1. Interview

Instructions :
**Exp 2 :** Hello ....... Please give me the notes you have taken and sit down opposite me. I'll now switch on the camera so that we can evaluate your behavior during the exercise. You can now explain your situation and the need for this pension. You have 5 minutes, use them.

**After 3 minutes (tick) :**

☐ If the participant talks for 3 minutes without stopping, interrupt her and ask her the following questions

☐ If the participant stops before 3 minutes, wait 20 seconds and then say “that's not enough, you still have time”. If she doesn't continue, add “there are missing elements to justify your claim”. If she still doesn't know what to say, ask her the questions below.

☐ After the 3-minute presentation, ask the patient a few questions (see questions below).

**Questions to ask the participant for 2 minutes, after the 3-minute speech (check those asked):**

☐ What do you do all day?

☐ What do you usually do to help with your pain?

☐ Is it possible to adapt or retrain your workstation?

☐ Do you think you'll be able to return to work? If so, how soon?

☐ What are you capable of doing?

☐ Why don't you want to work at this percentage?

☐ Why do you think we should allocate a pension to you?

☐ Have you really tried everything?

☐ How can you prove to me that you have this pathology? Do you have any medical tests to prove it?

☐ But do you actually want to work?

☐ This is insufficient to justify the requested pension.

☐ Please do an effort, it's not explained very well.

☐ I don’t believe you

☐ Give me a sentence that sums up your arguments: why you?

Instructions :

We've now completed our interview. Now I'm going to ask you to take the paper on your left and answer the 2 questions you've already seen.

### 2. Arithmetic task

I'd now like you to solve an arithmetic task as part of your assessment. Please count backwards from 2033 to 0, subtracting 13 each time and give me the results. Please calculate mentally as quickly and accurately as possible. If you make a calculation error, I'll inform you of your mistake and you'll have to start again from the beginning. Do you have any questions? Thank you, the time starts now.

Instructions for additional sentences :

Date : ___/___/___ Collaborator :

N° patient :

→ In the event of an error: “This is incorrect, please start again from 2033”.

→ If calculation aloud: “Please calculate the correct answer in your head and then give it to me aloud.”

→ If they refuse: “Try to do your best. In a moment, if you haven't been able to finish, we'll stop this task”. - if the participant clearly says she wants to stop, stop.

→ When the 1916 threshold is reached, “Please be faster in your calculations” to be repeated as many times as necessary = in bold (ok to recall where it was if asked).

□ 2033
□ 2020
□ 2007
□ 1994
□ 1981
□ 1968
□ 1955
□ 1942
□ 1929
□ **1916 a bit faster please »**
□ 1903
□ 1890
□ 1877
□ 1864
□ 1851

□ 1838
□ 1825
□ 1812
□ 1799
□ **1786**
□ 1773
□ 1760
□ 1747
□ 1734
□ 1721
□ 1708
□ 1695
□ 1682
□ 1669
□ **1656**
□ 1643

□ 1630
□ 1617
□ 1604
□ 1591
□ 1578
□ 1565
□ 1552
□ 1539
□ **1526**
□ 1513
□ 1500
□ 1487
□ 1474
□ 1461
□ 1448
□ 1435

□ 1422
□ 1409
□ **1396**
□ 1383
□ 1370
□ 1357
□ 1344
□ 1331
□ 1318
□ 1305
□ 1292
□ 1279
□ **1266**

Number of mistakes : ________________

Instructions :
Thank you very much, we've finished. (Opens door) Goodbye.

### Notes on the patient's attitude:

______________________________________________________________________

**How stressed does the patient appear during the interview?**

**Not at all** ◆━━━━━━━━━━━━━━━━━━◆ **A lot**

**How stressed does the patient appear during the arithmetic task?**

**Not at all** ◆━━━━━━━━━━━━━━━━━━◆ **A lot**

**How “mean” was I during the interview?**

**Not at all** ◆━━━━━━━━━━━━━━━━━━◆ **A lot**

**How “mean” was I during the arithmetic task ?**

**Not at all** ◆━━━━━━━━━━━━━━━━━━◆ **A lot**

Date : ___/___/___ Collaborator :
